# Towards a molecular systems model of coronary artery disease

**DOI:** 10.1101/003525

**Authors:** Gad Abraham, Oneil G. Bhalala, Paul I.W. de Bakker, Samuli Ripatti, Michael Inouye

## Abstract

Coronary artery disease (CAD) is a complex disease driven by myriad interactions of genetics and environmental factors. Traditionally, studies have analyzed only one disease factor at a time, providing useful but limited understanding of the underlying etiology. Recent advances in cost-effective and high-throughput technologies, such as single nucleotide polymorphism (SNP) genotyping, exome/genome sequencing, gene expression microarrays and metabolomics assays have enabled the collection of millions of data points in many thousands of individuals. In order to make sense of such ‘omics’ data, effective analytical methods are needed. We review and highlight some of the main results in this area, focusing on integrative approaches that consider multiple modalities simultaneously. Such analyses have the potential to uncover the genetic basis of CAD, produce genomic risk scores (GRS) for disease prediction, disentangle the complex interactions underlying disease, and predict response to treatment.

## Introduction

Coronary artery disease (CAD) is a significant impediment to good health and productivity worldwide. Globally, CAD is the leading cause of death with 7 million deaths in 2011 alone, accounting for 11.2% of all deaths [1]. Unfortunately the impact of CAD is increasing, as the disease burden is projected to nearly double from 47 million disability-adjusted life years (DALYs) in 1990 to 82 million DALYs in 2020 [2]. However, this impact is asymmetrically distributed between developed and developing countries. CAD morbidity is projected to more than double in developing countries from 1990 to 2020, but only increase by 50% in developed countries [3].

Development of CAD is a multi-decade process of atherosclerotic formation and chronic inflammation that ultimately leads to angina, myocardial infarction (MI), and death [4]. It arises from complex interactions of a multitude of factors - both environmental and genetic. Risk factors, such as tobacco use, low physical activity, obesity, hypertension, hypercholesterolemia, and diabetes have long been known. The contribution of genetics has been suspected given the importance of family history for CAD in an individual’s current risk. This has been strengthened with data from Swedish twin studies demonstrating a substantially larger concordance rate for monozygotic twins compared to dizygotic twins and heritability of 0.57 for males and 0.38 for females [5,6]. However, identifying the specific genetic changes that modify CAD risk has been and still is a challenge despite intense investigation in recent years.

## Uncovering the Genetic Basis of CAD

The publication of the Human Genome in 2001 offered an unprecedented launching pad for the understanding the genetic basis of diseases [7,8]. Within six years, seminal papers were published that began to outline the genetic architecture of CAD. These were initially based on univariate genome wide association studies (GWAS), where frequencies of individual SNPs in those with CAD (cases) were compared to those without CAD (controls). Although these GWAS datasets were significantly larger than what researchers were used to at that time, the statistical approach of testing individual SNPs for association was rather straightforward. One of the first studies considered 1,607 MI cases and 6,728 controls with no history of CAD from an Icelandic population [9]. This genome-wide scan found and replicated a locus associated with MI whose top SNP yielded an odds ratio of 1.28.

In the same issue of *Science*, McPherson et al also found a locus associated with CAD in the Ottawa Heart Study that validated in both the Copenhagen City Heart Study and Dallas Heart Study [10]. The top SNP in the locus increased the risk of CAD by 15-20% in individuals who were heterozygous and 30-40% in those who were homozygous. Analysis of CAD cases from the Wellcome Trust Case Control Consortium [11] and German MI Family Study revealed another SNP that was also strongly associated with CAD [12]. Each copy of the SNP allele increased CAD risk by 36%.

Interestingly, these initial SNPs all mapped to chromosome band 9p21, identifying it as the first locus to harbour common variants for CAD [13]. This locus was also replicated in other populations including Italian, Japanese, Korean, and Indian, suggesting it may be a universal risk allele [14–16]. 9p21 does not code for any known proteins but the closest genes, within 150kb, are *CDKN2A* and *CDKN2B*, which control cell proliferation and apoptosis. Moreover, targeted deletion of the non-coding interval of 9p21 in mice increased cardiac expression of *CDKN2A* and *CDKN2B* as well as vascular cell proliferation [17]. In addition, it has also been shown that 9p21 contains an abundance of enhancers and that the CAD risk SNPs disrupt STAT1 binding to one such enhancer thereby disrupting the interferon-γ response [18]. The mediator between 9p21 and downstream genes may be *ANRIL*, a non-coding RNA in the region which can alter downstream gene expression [19,20]. 9p21 has also been found to be associated with coronary calcification levels, abdominal aortic aneurysms and intracranial aneurysms, suggesting a broader role for this variant in vessel function [21–24].

CAD has been associated with other loci in subsequent large-scale meta-analyses. A variant mapping to 6q25.1 increases CAD risk by 23% per allele [12]. Notably, this SNP is located within the *MTHFD1L* gene, which can influence plasma homocysteine levels and affect risk of CAD [11]. New loci have also been identified using haplotype analysis, whereby proximal SNPs are grouped together and treated as a single unit, thus enhancing the detection power. Re-analysis of the WTCCC dataset identified the 6q26-q27 locus, containing four SNPs within the *SLC22A3-LPAL2-LPA* gene cluster [25]. The *LPA* gene encodes for part of lipoprotein(a) (Lp(a)), whose plasma levels correlate with CAD pathogenesis [26,27]. Analysis of the haplotype determined that it accounted for 15% of Lp(a) plasma level variability. Soranzo et al also used haplotype analysis to identify 12q24 as a CAD locus containing two SNPs yielding an odds ratio of 1.144 [28]. Interestingly, this locus also demonstrates considerably disease pleiotropy with type 1 diabetes, celiac disease and hypertension.

## “Missing” Heritability

The initial univariate GWAS were able to identify common SNPs associated with CAD with low to moderate effect size [29]. However, these SNPs account for less than 10% of CAD heritability, and it has been posited that the remaining unexplained heritability may be due to rarer variants with larger effects and/or undiscovered common variants with small effects [30–32]. These shortcomings suggested the need for larger data sets to allow for increased detection power. The CARDIoGRAM Consortium was formed to overcome these issues by amalgamating 14 GWAS with 22,223 CAD cases and 64,762 controls of European ancestry [33]. This meta-analysis identified 13 new loci, with each allele increasing CAD risk by between 6% to 17% [34]. The C4D Genetics Consortium performed another large-scale analysis, where 8,424 cases of European ancestry and 6,996 cases of South Asian ancestry, along with 15,062 controls were analysed [35]. Five new loci were identified including one SNP of particular biological interest as the nearest gene, *PDGFD* (117kb downstream), encodes the platelet-derived growth factor D protein. This protein is suspected to promote pathogenesis of atherosclerotic plaques and the SNP’s risk allele was associated with increased transcription of *PDGFD* in aortic media, aortic adventitia and mammary artery. Together, both CARDIoGRAM and C4D recently identified 46 genome-wide significant loci, explaining a little over 10% of CAD heritability [36]. While many CAD SNPs are still to be discovered, more collaborative GWAS with larger and more diverse sample populations are needed to further explain the remaining “missing” heritability.

## Constructing Genomic Risk Scores

One of the potential benefits of genomic association studies is the use of an individual’s genomic profile to create a score capturing the risk of developing CAD. Currently, the Framingham Risk Score (FRS) is a powerful tool that incorporates a patient’s age, sex, cholesterol, smoking status, blood pressure, and diabetes status to generate a 10-year CAD risk prediction [37]. However, it can be the case that a majority of new CAD cases appear in individuals who are not in the highest FRS risk group [38]. Given that many of the FRS factors are dependent on age and/or have a genetic component, the use of genetic variants may allow for earlier and more efficient identification of those at risk for CAD before changes in other risk factors like blood pressure or plasma lipid levels occur. Therefore, a genomic risk score (GRS), comprised of the increased risk associated with genetic variants, can be calculated and used to stratify patients into different risk categories, as is done with the FRS; early stratification being crucial for maximum benefit from lifestyle modification or therapeutic intervention to stymie pathogenesis.

As an initial investigation of the usefulness of a GRS with regard to individual risk prediction, Ripatti et al built a 13-SNP GRS for CAD and tested its utility in a prospective study of 30,725 individuals of European ancestry free of cardiovascular disease [39]. They found that patients with a GRS in the top 20% had a 66% increased risk (95% CI: 35-104%, P=10^-10^). In another study, a very similar 13-SNP GRS was tested in 3,014 individuals from the Framingham Heart Study with an 11-year median follow-up [40]. Analysis in this population found that each risk allele had a hazard ratio of 1.07 (95% CI: 1.00-1.15, P=0.04). While this GRS did not improve prediction of CAD over the FRS, it did associate strongly with high coronary artery calcium levels, a marker for subclinical atherosclerosis, suggesting that these SNPs may be predictive of atherosclerotic plaque progression.

In another prospective study of 4,818 Caucasian males [41], a 15-SNP GRS improved net reclassification (6.5%, P=0.044) and discrimination (1.11%, P=0.048) over FRS, indicating that there was a marginal improvement of identifying those at risk for CAD [42]. The largest benefit of the 15-SNP GRS was realized when analysis was restricted to men between 50 and 59 years, where a 2.8% (P=0.0038) discrimination improvement over FRS occurred. Analysis of a 28-SNP GRS in a Finnish-based prospective cohort of 24,124 individuals also showed an improvement over FRS [43]. In this study, FRS provided a discrimination index of 0.851, while adding the GRS improved the index to 0.856 (P=0.0002). Moreover, addition of the GRS helped reclassify 12% of the individuals as high-risk, potentially identifying a group of patients that would benefit from early targeted healthcare. The value of a GRS also extends beyond individual risk prediction as it also captures a “best guess” of the genetic architecture of disease/trait, thus genetic overlaps between CAD traits can also be assessed [44].

While it is still early days, these GRS studies point to the potential use of SNPs in identifying and classifying individuals as high- and low-risk for CAD. It is also clear that GRS’s based on lead SNPs from each associated locus use only a small fraction of the potential for prediction and more elaborate strategies and statistical methods, such as supervised learning, are needed to optimize the genomic predictions [45]. Further analyses using large numbers of genetic variants are needed to determine the effectiveness of a GRS as a screening tool and indicator for intervention, especially in or before the early stages of pathogenesis where FRS variables like age, cholesterol and blood pressure are less predictive and lifestyle modifications can have a greater impact.

## Network Approaches to Understand CAD Etiology

Despite the success of GWAS, these results only provide one piece of the puzzle. It has been challenging to interpret the individual SNP associations in terms of the underlying biological pathways, and therefore current knowledge of disease etiology is still limited. To have a phenotypic effect, true causal genetic variants must exert their influence through other mediators, such as changes in the structure of protein encoded by the gene (e.g. non-synonymous variants) or changes in regulatory elements affecting the binding affinity of transcription factors, thus affecting the gene’s RNA levels and presumably the levels of the encoded protein. Data about these biological phenomena were not available in many early studies, limiting the ability to decipher these complex effects. To address this shortcoming, studies now aim to combine various sources of systems level information, such as genomic, transcriptomic, and metabolomic data. Since both gene expression and metabolites are known to be influenced by genetic variation, joint investigation of these genetic effects on gene expression and metabolite levels and on clinically-relevant phenotypes such as atherosclerosis affords insight into how the genetic effects on disease are mediated [46].

Beyond the ability to examine multiple sources of biological data at once, another important conceptual advance has been the shift towards examining genes as part of networks, such as transcriptional regulation networks [47], rather than considering them in isolation. A network-based analysis has the power to better model these biomolecular relationships and their potential role in disease. The network is the context: in some cases, the individual gene associations may be weak but subtle global trends may reveal themselves when examined at the level of multiple related genes.

In a formal mathematical sense, a network consists of *nodes* (also called *vertices*) connected by *edges* (see Box 1). Typically, nodes represent some discrete entity, such as a gene, protein or metabolite, whereas edges represent relationships between entities. The edges may be *undirected*, representing a symmetric relationship such as “gene A is correlated with gene B” or *directed*, representing an asymmetric relationship such as “gene A up-regulates gene B”. The edges need not represent purely physical relationships such as interactions; they may simply represent statistical effects such as correlation. One of the earliest and simplest applications in network analysis has been visualization of complex relationships between biological entities such as genes and genetic variants, allowing the researcher to construct a mental image of the underlying etiology. However, as the data becomes larger, more complex, and multi-faceted so the networks become less interpretable, requiring more sophisticated approaches for visualization and analysis in order to extract biological meaning from them.

Researchers have long known that information sources such as transcriptomics can offer valuable insights into CAD etiology [48,49], as circulating leukocytes link organ systems in mediating the atherosclerotic process [50], however integration with other data sources including genetic and metabolite variation has only been possible relatively recently with the wide-spread use of high-throughput assays. One of the first examples of network-driven models of complex disease is that by Chen et al. [51], who examined genotypes, gene expression, and metabolite data from BxH-cross mice. They constructed networks of co-expressed genes and identified highly-connected components (modules) of these networks that were associated both with genetic variants and with several intermediate phenotypes related to obesity, diabetes and atherosclerosis, such as abdominal fat mass, weight, plasma insulin, free fatty acids, total plasma cholesterol and aortic lesion size. Using a statistical approach based on Mendelian Randomization [52] (see below), some of these sub-networks were then postulated as causal mediators between genetic variants and the disease-related traits. One of the modules with the strongest associations that was also supported to have a causal effect on metabolic traits was the macrophage-enriched metabolic network (MEMN) consisting of 1,406 genes, out of which 375 genes were estimated to be causal of obesity-related traits. From these genes, it was hypothesized that *Lpl* and *Lactb* caused obesity and that *Ppm1l* was a causal driver of phenotypes related to metabolic syndrome, predictions which were subsequently validated in mice [53]. Significantly, the MEMN network together with its link to obesity were also replicated in human adipose tissue [54]. Beyond the specific findings of these studies, these results highlight the power of integrating several sources of information and of considering networks rather than genes in isolation to uncover some of the biological processes connecting genetic variation with observable phenotypes.

## Mendelian Randomization for Predicting Causality

Network analyses tend to rely on statistical association, such as correlation, to infer the network structure from observational data. In order to identify new drug targets and interventions that can reduce disease risk, we must identify which factors cause disease and which are merely associated with it, as association does not necessarily imply causation [55]. For example, if a metabolite is observed to be associated with disease, one cannot say that the metabolite is causal of the disease based on this observation alone. Such a claim can only be made using methods such as interventional experiments where the metabolite level is manipulated and the resulting effect on disease is observed. Yet, in some cases, statistical analyses of genomic variation may be able to determine which relationships are causal. Genetic variation is unique in that an individual’s alleles are assigned randomly during meiosis (assuming non-assortative mating with respect to the phenotype of interest) and are largely fixed for the life of the individual. Consequently, genetic variants can be viewed as natural perturbations of the system (Mendelian Randomization [56,57]), allowing us to interpret a genetic-phenotypic association as causal in one direction, namely genetic variant causes phenotype. Leveraging this causal information through statistical frameworks such as instrumental variables (IVs) allows one to infer whether other associations with disease are indeed causal or simply correlations. Such an approach has provided evidence that increases in plasma HDL cholesterol do not lower the risk of myocardial infarction despite the well-documented strong negative correlation between the two in observational studies, thus drug targeting of HDL-C for CAD treatment may not be a successful strategy [58]. Applied on a larger scale, statistical methods based on the principles of Mendelian Randomization approaches can help orient the edges in an undirected gene or metabolite network (inferred from associations), generating directed networks representing putative causal structures [52,59,60] that can later be tested experimentally either in laboratory models or randomized controlled trials in humans.

## Integrative Omics to Decipher the Role of Inflammation in CAD

In humans, large-scale multi-omic datasets are being increasingly utilized to better elucidate the biological pathways responsible for the known links between inflammation and CAD [61]. Laurila et al. [62] analyzed Finnish individuals assayed for transcriptomics in fat tissue, HDL lipidomic profiles, and genotypes, in order to dissect the genetic contribution to levels of plasma high-density lipoprotein cholesterol (HDL-C), as HDL-C levels are known to be negatively associated with risk of CAD. The analysis compared multi-omic profiles of individuals at the extremes of HDL distribution and highlighted the role of the *HLA* region and of inflammation pathways in controlling HDL-C levels and a wide range of differences in both adipose transcriptome and lipidomic profiles. Another analysis of CAD case/control individuals from the Framingham Heart Study [63] revealed differences in co-expression patterns of genes between cases and controls; these *differential modules* were found to be enriched for quantitative trait loci (QTLs) that affect CAD risk, including a module enriched for B-cell immune genes. By integrating various data sources, including protein-protein interaction (PPI) networks and gene networks derived from other studies, they found several regulatory genes, dubbed *key drivers*, exhibiting strong effects on these modules. One such gene was *TNFRSF13C*, which is known to affect aortic root atherosclerosis, thereby providing support for the hypothesis that changes in co-regulation of B-cell-related gene networks, caused by such drivers, was partly responsible for increases in CAD risk.

Expanding beyond the small set of metabolites traditionally associated with CAD (e.g. total LDL/HDL cholesterol or triglycerides), characterization of a wide-range of metabolite levels has become increasingly important in understanding of etiology CAD and related metabolic diseases [64,65]. Atherosclerotic plaques themselves are heterogeneous and composed of multiple immune cell types and a wide variety of fats, lipoproteins, and other metabolites [66,67]. Thus there is a need to include metabolomic variation together with genetic and transcriptomic effects. Such analyses have now become possible with technological advances in ^1^H NMR and high-resolution mass spectrometry, which routinely measure levels of hundreds of metabolite species in large human cohorts. For example, a recent metabolomic GWAS of 216 serum metabolite measures in over 8,300 Finnish individuals [68] identified 31 loci with genome-wide significant associations to metabolite levels. Integration of genomics, transcriptomic and metabolite data in a large Finnish cohort led to specific evidence for the role of inflammation in CAD through the identification of the Lipid-Leukocyte (LL) module [69], a network of highly co-expressed genes related to the acute inflammatory response, which was inferred to be reactive to lipid levels through causal inference methods [59]. This analysis was later extended to assess a wide-range of metabolite species, revealing the effect of specific sub-species on the module’s coherence, indicating that the degree of co-expression in the module was significantly associated with metabolite levels such as linoleic acid and various LDL and HDL particles [70]. Later, by integrating serum metabolomics data with genetic variation and transcriptomic data in humans and in mice, Inouye et al. [46] used a powerful data-driven multivariate approach to identify 11 metabolic networks and detect 7 previously-unknown loci associated with serum metabolite levels, notably *SERPINA1* and *AQP9*. Transcriptional data was then used to show that *AQP9* expression in murine liver was associated with the size of atherosclerotic lesion and, in humans, both *AQP9* and *SERPINA1* expression in arterial tissue was substantially up-regulated in plaques, providing a potential explanatory link between genetic variation and a phenotype relevant to disease progression.

## New Data Sources to Model Disease

Apart from genetic, transcriptomic, and metabolomic data, another rich source of information is the microbiome, the characterization of the diverse set of microbial communities inhabiting the human body, which are increasingly being recognized as important factors in obesity and atherosclerosis [71–73]. Recent evidence points to the presence of oral pathogens such as *Chryseomonas, Veillonella*, and *Streptococcus* in atherosclerotic plaques and thus potentially contributing to the inflammatory process [74,75]. A recent example of one such integrative study involving the gut microbiome, genetic variation, and gene expression in inbred mice strains (Hybrid Mouse Diversity Panel) [76] across several time points revealed the complex interactions between these components in contributing to obesity. Further, the effects of high-fat/high-sucrose diets on the composition of gut bacterial communities was modulated by the specific genetic makeup of each mouse strain, leading to down-stream effects on metabolism and eventual risk of obesity.

Another potential source of useful information is epigenetics, which describes both epigenetic marks and non-coding RNAs (ncRNAs) [77,78]. Many epigenetic marks are thought to be reset during early embryogenesis; however, some epigenetic marks may still be passed between generations [79], and marks may be modified in response to environmental exposures such as smoking [80]. Some epigenetic effects relevant to CAD include homocysteine-induced methylation in vascular smooth muscle cells, contributing to atherosclerosis [81], and the ncRNA microRNA-33 found to regulate cholesterol homeostasis [82]. Further large-scale studies will be required in order to assay genome-wide methylation status and ncRNA expression levels in concert with other data sources.

## Future Directions

Apart from incorporating novel sources of biological variation, one important factor not considered in many studies is time itself: the biological processes underlying cardiac and metabolic disease are dynamic, all while interacting with a complex array of environmental and genetic effects. CAD may take several decades to manifest clinically, and is typically preceded by sub-clinical phenomena. Clearly, this disease progression can be influenced at various stages by external interventions such as life-style changes (diet and exercise) and medication such as statins, and disease trajectories vary widely between individuals. Hence, to better understand these processes larger and more detailed repeated measures data will be required. Such experiments present their own unique challenges in analysis and interpretation [83,84]. A recent multi-omic study examined whole-genome sequencing, RNA sequencing, proteomics, metabolomics, ncRNA, and auto-antibody data measured in blood components from one individual over a 14 month period [85], revealing in detail the correlation of multiple body systems like inflammatory and insulin response pathways to viral infection and early onset of type 2 diabetes. This study provides a glimpse into what will become more common, with costs of technology reducing to a level enabling such detailed measurements over much larger cohorts and longer time-scales. Similar studies with much larger sample sizes based on individuals with various disease states will be necessary to be able to draw robust conclusions about the pathogenesis of CAD and improve preclinical models.

To complement temporal dynamics, spatial effects will need to be considered as well, that is, heterogeneity in expression across tissues [86,87]. Studies that consider only one tissue type, typically blood and its components, will not be able to detect such variation. This will require assays of multiple tissue types, both to capture coordinated processes happening in all tissues and to localize effects that occur only in specific tissues. One such effort is the Genotype-Tissue Expression (GTEx) project [88], which aims to conduct a wide-ranging survey of gene expression and its associated genetic variation across multiple human tissues.

In addition to the important goals of predicting those individuals that are at high risk or disease and developing a deeper understanding of disease etiology, another major avenue of research where systems and network approaches may have an impact is pharmacogenomics, that is, the study of how genetic variation influences each individual’s response to medication. Both drug response and drug metabolism can be considered as phenotypes, and many of the existing analysis methods are applicable. This will be useful for tailoring more specific sets of medication to each individual, matched to their genomic profile [89,90]. The clinical utility of pharmacogenomics approaches has so far been rather limited [91]. However, testing for loss-of-function variation in the gene *CYP2C19* (part of the P450 family of enzymes) is now recommended for informing the use of the antiplatelet drug clopidogrel in individuals with acute coronary syndrome (ACS) undergoing percutaneous coronary intervention (PCI) [92], since such variants may adversely affect platelet activity and increase the risk of cardiovascular events.

## Conclusions

CAD and related diseases are complex phenotypes and are the result of interplay amongst a multitude of genetic and environmental effects together with an important inflammatory aspect. Initially, analyses tended to be restricted to only one type of high throughput data, such as GWAS considering association of disease with genetic markers, or only considering blood biomarkers for disease progression. While such focused analyses have proven insightful, they provide limited information about a narrow aspect of the disease, while ignoring the complex interactions between these different processes. The advent of multi-omic studies which concurrently analyze genetic variation, transcriptomics, metabolomics and others sources of information over hundreds or thousands of individuals, has begun providing deeper insight into the underlying mechanisms responsible for observed phenomena, such as the gene networks responsible for the known link between inflammation and disease and the various feedback loops amongst genetic variation, microbial communities, metabolism, and ultimately disease.

Hence, the coming major challenges will firstly be collection of large-scale multi-omic samples across thousands of individuals in order to achieve the statistical power to detect the multitude of interacting effects in CAD etiology. Secondly, novel statistical and computational methods will be required to effectively combine these disparate information sources into a coherent model of disease, with the aim of generating plausible biological hypotheses that can be tested in animal models and clinical trials. Thirdly, the testing of these models will need to rigorously evaluate the specific reliability of the models as well as their modes of failure so that refinements to the model can be made. Meeting these challenges will result in greater understanding of CAD etiology and, if successfully implemented, translate into better clinical outcomes.

**Figure 1.**
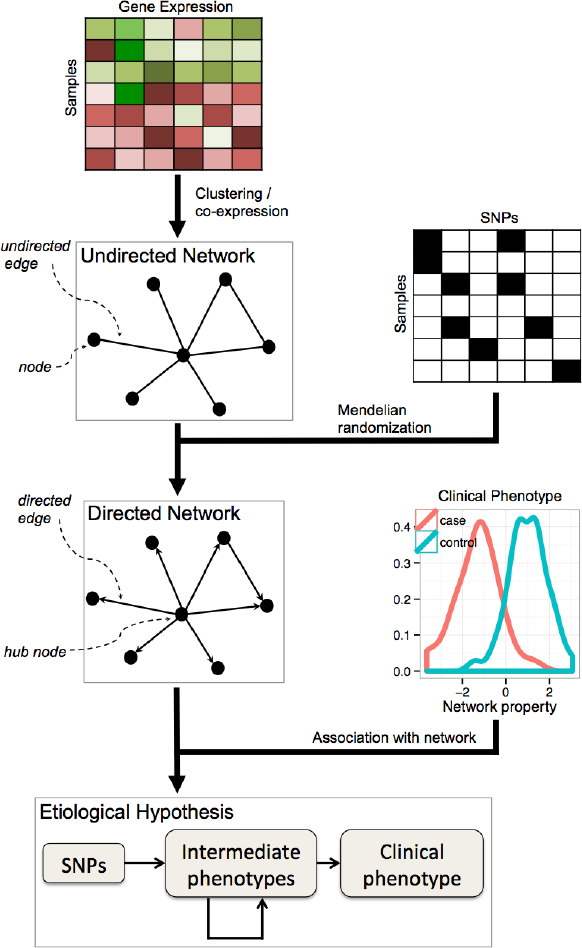
Network-based analysis of omic data to model the processes connecting genetic variation to disease.

**Table 1:** Molecular networks relevant to coronary artery disease

| Name | Source | Main associations | Biological interpretation | Ref |
| --- | --- | --- | --- | --- |
| **Macrophage-enriched metabolic network (MEMN)** | B×H mouse cross, multiple tissues | Obesity, diabetes, atherosclerosis, total cholesterol levels, and HDL levels | Network represents an inflammatory response driven by genetic variation and affecting disease phenotypes, in liver and/or adipose tissue. | [51] |
| **Lipid-leukocyte (LL) module** | Human population-based cohort, whole blood | Various metabolites (HDL, VLDL, glycoproteins, isoleucine, and others) and immune response markers (IL-1ra, CRP, and heavy molecular weight adiponectin) | Module implicates acute inflammatory cells (mast cells and basophils) as reactive to various metabolites. Inverse relationship between module expression and its density. | [69,70] |
| **Combined Inflammatory Pathway** | Human cohort enriched for low familial HDL-C, adipose tissue | VCAM1 levels and SNPs predictive of low HDL-C | Pathway represents an inflammatory link between genetic variation and HDL-C levels and indicates that HDL levels may be regulated by inflammatory processes. | [62] |
| **Two differential modules (case and control)** | Human CAD case/control study, whole blood | Control module enriched for SNPs predictive of CAD. | Differential pathways represent two modes of regulation acting in CAD vs. non-CAD individuals, and indicate that B-cell immune pathways may be causally driving lipids levels and risk of CAD. | [63] |

HDL: high density lipoprotein
VLDL: very low density lipoprotein
IgE: immunoglobulin E
IL-1ra: Interleukin 1 receptor antagonist
CRP: C-reactive protein
HMWA: high molecular weight adiponectin
VCAM1: vascular cell adhesion molecule 1

## References

1. Organization WH (2013) The top 10 causes of death: Fact Sheet.

2. Mackay J, Mensah GA, Mendis S, Greenlund K, World Health Organization. (2004) The atlas of heart disease and stroke. Geneva: World Health Organization. 112 p. p.

3. Okrainec K, Banerjee DK, Eisenberg MJ (2004) Coronary artery disease in the developing world. Am Heart J 148: 7–15.

4. Ross R (1999) Atherosclerosis—an inflammatory disease. N Engl J Med 340: 115–126.

5. Zdravkovic S, Wienke A, Pedersen NL, Marenberg ME, Yashin AI, et al. (2002) Heritability of death from coronary heart disease: a 36-year follow-up of 20 966 Swedish twins. J Intern Med 252: 247–254.

6. Marenberg ME, Risch N, Berkman LF, Floderus B, de Faire U (1994) Genetic susceptibility to death from coronary heart disease in a study of twins. N Engl J Med 330: 1041–1046.

7. Lander ES, Linton LM, Birren B, Nusbaum C, Zody MC, et al. (2001) Initial sequencing and analysis of the human genome. Nature 409: 860–921.

8. Venter JC, Adams MD, Myers EW, Li PW, Mural RJ, et al. (2001) The sequence of the human genome. Science 291: 1304–1351.

9. Helgadottir A, Thorleifsson G, Manolescu A, Gretarsdottir S, Blondal T, et al. (2007) A common variant on chromosome 9p21 affects the risk of myocardial infarction. Science 316: 1491–1493.

10. McPherson R, Pertsemlidis A, Kavaslar N, Stewart A, Roberts R, et al. (2007) A common allele on chromosome 9 associated with coronary heart disease. Science 316: 1488–1491.

11. Wellcome Trust Case Control C (2007) Genome-wide association study of 14,000 cases of seven common diseases and 3,000 shared controls. Nature 447: 661–678.

12. Samani NJ, Erdmann J, Hall AS, Hengstenberg C, Mangino M, et al. (2007) Genomewide association analysis of coronary artery disease. N Engl J Med 357: 443–453.

13. Roberts R, Stewart AF (2012) Genetics of coronary artery disease in the 21st century. Clin Cardiol 35: 536–540.

14. Shen GQ, Rao S, Martinelli N, Li L, Olivieri O, et al. (2008) Association between four SNPs on chromosome 9p21 and myocardial infarction is replicated in an Italian population. J Hum Genet 53: 144–150.

15. Hinohara K, Nakajima T, Takahashi M, Hohda S, Sasaoka T, et al. (2008) Replication of the association between a chromosome 9p21 polymorphism and coronary artery disease in Japanese and Korean populations. J Hum Genet 53: 357–359.

16. Maitra A, Dash D, John S, Sannappa PR, Das AP, et al. (2009) A common variant in chromosome 9p21 associated with coronary artery disease in Asian Indians. J Genet 88: 113–118.

17. Visel A, Zhu Y, May D, Afzal V, Gong E, et al. (2010) Targeted deletion of the 9p21 non-coding coronary artery disease risk interval in mice. Nature 464: 409–412.

18. Harismendy O, Notani D, Song X, Rahim NG, Tanasa B, et al. (2011) 9p21 DNA variants associated with coronary artery disease impair interferon-gamma signalling response. Nature 470: 264–268.

19. Cunnington MS, Santibanez Koref M, Mayosi BM, Burn J, Keavney B (2010) Chromosome 9p21 SNPs Associated with Multiple Disease Phenotypes Correlate with ANRIL Expression. PLoS Genet 6: e1000899.

20. Liu Y, Sanoff HK, Cho H, Burd CE, Torrice C, et al. (2009) INK4/ARF transcript expression is associated with chromosome 9p21 variants linked to atherosclerosis. PLoS One 4: e5027.

21. O’Donnell CJ, Kavousi M, Smith AV, Kardia SL, Feitosa MF, et al. (2011) Genome-wide association study for coronary artery calcification with follow-up in myocardial infarction. Circulation 124: 2855–2864.

22. van Setten J, Isgum I, Smolonska J, Ripke S, de Jong PA, et al. (2013) Genome-wide association study of coronary and aortic calcification implicates risk loci for coronary artery disease and myocardial infarction. Atherosclerosis 228: 400–405.

23. Shea J, Agarwala V, Philippakis AA, Maguire J, Banks E, et al. (2011) Comparing strategies to fine-map the association of common SNPs at chromosome 9p21 with type 2 diabetes and myocardial infarction. Nat Genet 43: 801–805.

24. Helgadottir A, Thorleifsson G, Magnusson KP, Gretarsdottir S, Steinthorsdottir V, et al. (2008) The same sequence variant on 9p21 associates with myocardial infarction, abdominal aortic aneurysm and intracranial aneurysm. Nat Genet 40: 217–224.

25. Tregouet DA, Konig IR, Erdmann J, Munteanu A, Braund PS, et al. (2009) Genome-wide haplotype association study identifies the SLC22A3-LPAL2-LPA gene cluster as a risk locus for coronary artery disease. Nat Genet 41: 283–285.

26. Brazier L, Tiret L, Luc G, Arveiler D, Ruidavets JB, et al. (1999) Sequence polymorphisms in the apolipoprotein(a) gene and their association with lipoprotein(a) levels and myocardial infarction. The ECTIM Study. Atherosclerosis 144: 323–333.

27. Keenan TE, Rader DJ (2013) Genetics of lipid traits and relationship to coronary artery disease. Curr Cardiol Rep 15: 396.

28. Soranzo N, Spector TD, Mangino M, Kuhnel B, Rendon A, et al. (2009) A genome-wide meta-analysis identifies 22 loci associated with eight hematological parameters in the HaemGen consortium. Nat Genet 41: 1182–1190.

29. McPherson R (2013) From genome-wide association studies to functional genomics: new insights into cardiovascular disease. Can J Cardiol 29: 23–29.

30. Schunkert H, Erdmann J, Samani NJ (2010) Genetics of myocardial infarction: a progress report. Eur Heart J 31: 918–925.

31. Manolio TA, Collins FS, Cox NJ, Goldstein DB, Hindorff LA, et al. (2009) Finding the missing heritability of complex diseases. Nature 461: 747–753.

32. Eichler EE, Flint J, Gibson G, Kong A, Leal SM, et al. (2010) Missing heritability and strategies for finding the underlying causes of complex disease. Nat Rev Genet 11: 446–450.

33. Preuss M, Konig IR, Thompson JR, Erdmann J, Absher D, et al. (2010) Design of the Coronary ARtery DIsease Genome-Wide Replication And Meta-Analysis (CARDIoGRAM) Study: A Genome-wide association meta-analysis involving more than 22 000 cases and 60 000 controls. Circ Cardiovasc Genet 3: 475–483.

34. Schunkert H, Konig IR, Kathiresan S, Reilly MP, Assimes TL, et al. (2011) Large-scale association analysis identifies 13 new susceptibility loci for coronary artery disease. Nat Genet 43: 333–338.

35. Coronary Artery Disease Genetics C (2011) A genome-wide association study in Europeans and South Asians identifies five new loci for coronary artery disease. Nat Genet 43: 339–344.

36. Deloukas P, Kanoni S, Willenborg C, Farrall M, Assimes TL, et al. (2013) Large-scale association analysis identifies new risk loci for coronary artery disease. Nat Genet 45: 25–33.

37. Wilson PW, D'Agostino RB, Levy D, Belanger AM, Silbershatz H, et al. (1998) Prediction of coronary heart disease using risk factor categories. Circulation 97: 1837–1847.

38. Ripatti SS, V. (2013) How could use of genetic markers prevent coronary heart disease events? Personalized Medicine 10: 769–771.

39. Ripatti S, Tikkanen E, Orho-Melander M, Havulinna AS, Silander K, et al. (2010) A multilocus genetic risk score for coronary heart disease: case-control and prospective cohort analyses. Lancet 376: 1393–1400.

40. Thanassoulis G, Peloso GM, Pencina MJ, Hoffmann U, Fox CS, et al. (2012) A genetic risk score is associated with incident cardiovascular disease and coronary artery calcium: the Framingham Heart Study. Circ Cardiovasc Genet 5: 113–121.

41. Evans A, Salomaa V, Kulathinal S, Asplund K, Cambien F, et al. (2005) MORGAM (an international pooling of cardiovascular cohorts). Int J Epidemiol 34: 21–27.

42. Hughes MF, Saarela O, Stritzke J, Kee F, Silander K, et al. (2012) Genetic markers enhance coronary risk prediction in men: the MORGAM prospective cohorts. PLoS One 7: e40922.

43. Tikkanen E, Havulinna AS, Palotie A, Salomaa V, Ripatti S (2013) Genetic risk prediction and a 2-stage risk screening strategy for coronary heart disease. Arterioscler Thromb Vasc Biol 33: 2261–2266.

44. van ’t Hof FN, Ruigrok YM, Baas AF, Kiemeney LA, Vermeulen SH, et al. (2013) Impact of inherited genetic variants associated with lipid profile, hypertension, and coronary artery disease on the risk of intracranial and abdominal aortic aneurysms. Circ Cardiovasc Genet 6: 264–270.

45. Abraham G, Kowalczyk A, Zobel J, Inouye M (2013) Performance and robustness of penalized and unpenalized methods for genetic prediction of complex human disease. Genet Epidemiol 37: 184–195.

46. Inouye M, Ripatti S, Kettunen J, Lyytikäinen L-P, Oksala N, et al. (2012) Novel Loci for Metabolic Networks and Multi-Tissue Expression Studies Reveal Genes for Atherosclerosis. PLoS Genetics 8: e1002907.

47. Alon U (2007) An introduction to systems biology : design principles of biological circuits. Boca Raton, FL: Chapman & Hall/CRC. xvi, 301 p., 304 p. of plates p.

48. Patino WD, Mian OY, Kang JG, Matoba S, Bartlett LD, et al. (2005) Circulating transcriptome reveals markers of atherosclerosis. Proc Natl Acad Sci U S A 102: 3423–3428.

49. Kleemann R, Verschuren L, van Erk MJ, Nikolsky Y, Cnubben NH, et al. (2007) Atherosclerosis and liver inflammation induced by increased dietary cholesterol intake: a combined transcriptomics and metabolomics analysis. Genome Biol 8: R200.

50. Swirski FK, Nahrendorf M (2013) Leukocyte behavior in atherosclerosis, myocardial infarction, and heart failure. Science 339: 161–166.

51. Chen Y, Zhu J, Lum PY, Yang X, Pinto S, et al. (2008) Variations in DNA elucidate molecular networks that cause disease. Nature 452: 429–435.

52. Schadt EE, Lamb J, Yang X, Zhu J, Edwards S, et al. (2005) An integrative genomics approach to infer causal associations between gene expression and disease. Nat Genet 37: 710–717.

53. Yang X, Deignan JL, Qi H, Zhu J, Qian S, et al. (2009) Validation of candidate causal genes for obesity that affect shared metabolic pathways and networks. Nat Genet 41: 415–423.

54. Emilsson V, Thorleifsson G, Zhang B, Leonardson AS, Zink F, et al. (2008) Genetics of gene expression and its effect on disease. Nature 452: 423–428.

55. Pearl J (2000) Causality : models, reasoning, and inference. Cambridge, U.K.; New York: Cambridge University Press. xvi, 384 p. p.

56. Lawlor DA, Harbord RM, Sterne JA, Timpson N, Davey Smith G (2008) Mendelian randomization: using genes as instruments for making causal inferences in epidemiology. Stat Med 27: 1133–1163.

57. Sheehan NA, Didelez V, Burton PR, Tobin MD (2008) Mendelian randomisation and causal inference in observational epidemiology. PLoS Med 5: e177.

58. Voight BF, Peloso GM, Orho-Melander M, Frikke-Schmidt R, Barbalic M, et al. (2012) Plasma HDL cholesterol and risk of myocardial infarction: a mendelian randomisation study. Lancet 380: 572–580.

59. Aten JE, Fuller TF, Lusis AJ, Horvath S (2008) Using genetic markers to orient the edges in quantitative trait networks: the NEO software. BMC Syst Biol 2: 34.

60. Logsdon BA, Mezey J (2010) Gene expression network reconstruction by convex feature selection when incorporating genetic perturbations. PLoS Comput Biol 6: e1001014.

61. Libby P, Lichtman AH, Hansson GK (2013) Immune effector mechanisms implicated in atherosclerosis: from mice to humans. Immunity 38: 1092–1104.

62. Laurila PP, Surakka I, Sarin AP, Yetukuri L, Hyotylainen T, et al. (2013) Genomic, transcriptomic, and lipidomic profiling highlights the role of inflammation in individuals with low high-density lipoprotein cholesterol. Arterioscler Thromb Vasc Biol 33: 847–857.

63. Huan T, Zhang B, Wang Z, Joehanes R, Zhu J, et al. (2013) A systems biology framework identifies molecular underpinnings of coronary heart disease. Arterioscler Thromb Vasc Biol 33: 1427–1434.

64. Suhre K, Gieger C (2012) Genetic variation in metabolic phenotypes: study designs and applications. Nature Reviews Genetics 13: 759–769.

65. Suhre K, Shin S-Y, Petersen A-K, Mohney RP, Meredith D, et al. (2011) Human metabolic individuality in biomedical and pharmaceutical research. Nature 477: 54–60.

66. Hansson GK, Jonasson L (2009) The discovery of cellular immunity in the atherosclerotic plaque. Arterioscler Thromb Vasc Biol 29: 1714–1717.

67. Insull W, Jr. (2009) The pathology of atherosclerosis: plaque development and plaque responses to medical treatment. Am J Med 122: S3–S14.

68. Kettunen J, Tukiainen T, Sarin A-P, Ortega-Alonso A, Tikkanen E, et al. (2012) Genome-wide association study identifies multiple loci influencing human serum metabolite levels. Nature Genetics 44: 269–276.

69. Tnouye M, Silander K, Hamalainen E, Salomaa V, Harald K, et al. (2010) An immune response network associated with blood lipid levels. PLoS Genet 6: e1001113.

70. Inouye M, Kettunen J, Soininen P, Silander K, Ripatti S, et al. (2010) Metabonomic, transcriptomic, and genomic variation of a population cohort. Molecular Systems Biology 6: 441.

71. Caesar R, Fak F, Backhed F (2010) Effects of gut microbiota on obesity and atherosclerosis via modulation of inflammation and lipid metabolism. J Intern Med 268: 320–328.

72. Turnbaugh PJ, Hamady M, Yatsunenko T, Cantarel BL, Duncan A, et al. (2009) A core gut microbiome in obese and lean twins. Nature 457: 480–484.

73. Le Chatelier E, Nielsen T, Qin J, Prifti E, Hildebrand F, et al. (2013) Richness of human gut microbiome correlates with metabolic markers. Nature 500: 541–546.

74. Gaetti-Jardim E, Jr., Marcelino SL, Feitosa AC, Romito GA, Avila-Campos MJ (2009) Quantitative detection of periodontopathic bacteria in atherosclerotic plaques from coronary arteries. J Med Microbiol 58: 1568–1575.

75. Koren O, Spor A, Felin J, Fak F, Stombaugh J, et al. (2011) Human oral, gut, and plaque microbiota in patients with atherosclerosis. Proc Natl Acad Sci U S A 108 Suppl 1: 4592–4598.

76. Parks BW, Nam E, Org E, Kostem E, Norheim F, et al. (2013) Genetic control of obesity and gut microbiota composition in response to high-fat, high-sucrose diet in mice. Cell Metab 17: 141–152.

77. Ordovas JM, Smith CE (2010) Epigenetics and cardiovascular disease. Nat Rev Cardiol 7: 510–519.

78. Gluckman PD, Hanson MA, Buklijas T, Low FM, Beedle AS (2009) Epigenetic mechanisms that underpin metabolic and cardiovascular diseases. Nat Rev Endocrinol 5: 401–408.

79. Hackett JA, Sengupta R, Zylicz JJ, Murakami K, Lee C, et al. (2013) Germline DNA demethylation dynamics and imprint erasure through 5-hydroxymethylcytosine. Science 339: 448–452.

80. Oka D, Yamashita S, Tomioka T, Nakanishi Y, Kato H, et al. (2009) The presence of aberrant DNA methylation in noncancerous esophageal mucosae in association with smoking history: a target for risk diagnosis and prevention of esophageal cancers. Cancer 115: 3412–3426.

81. Yideng J, Jianzhong Z, Ying H, Juan S, Jinge Z, et al. (2007) Homocysteine-mediated expression of SAHH, DNMTs, MBD2, and DNA hypomethylation potential pathogenic mechanism in VSMCs. DNA Cell Biol 26: 603–611.

82. Rayner KJ, Suarez Y, Davalos A, Parathath S, Fitzgerald ML, et al. (2010) MiR-33 contributes to the regulation of cholesterol homeostasis. Science 328: 1570–1573.

83. Yeung KY, Dombek KM, Lo K, Mittler JE, Zhu J, et al. (2011) Construction of regulatory networks using expression time-series data of a genotyped population. Proc Natl Acad Sci U S A 108: 19436–19441.

84. Lebre S, Becq J, Devaux F, Stumpf MP, Lelandais G (2010) Statistical inference of the time-varying structure of gene-regulation networks. BMC Syst Biol 4: 130.

85. Chen R, Mias GI, Li-Pook-Than J, Jiang L, Lam HY, et al. (2012) Personal omics profiling reveals dynamic molecular and medical phenotypes. Cell 148: 1293–1307.

86. Nica AC, Parts L, Glass D, Nisbet J, Barrett A, et al. (2011) The architecture of gene regulatory variation across multiple human tissues: the MuTHER study. PLoS Genet 7: e1002003.

87. Dimas AS, Deutsch S, Stranger BE, Montgomery SB, Borel C, et al. (2009) Common regulatory variation impacts gene expression in a cell type-dependent manner. Science 325: 1246–1250.

88. GTEx Consortium (2013) The Genotype-Tissue Expression (GTEx) project. Nat Genet 45: 580–585.

89. Roden DM, Johnson JA, Kimmel SE, Krauss RM, Medina MW, et al. (2011) Cardiovascular pharmacogenomics. Circ Res 109: 807–820.

90. Weeke P, Roden DM (2013) Pharmacogenomics and cardiovascular disease. Curr Cardiol Rep 15: 376.

91. Ioannidis JP (2013) To replicate or not to replicate: the case of pharmacogenetic studies: Have pharmacogenomics failed, or do they just need larger-scale evidence and more replication? Circ Cardiovasc Genet 6: 413–418; discussion 418.

92. Scott SA, Sangkuhl K, Stein CM, Hulot JS, Mega JL, et al. (2013) Clinical pharmacogenetics implementation consortium guidelines for CYP2C19 genotype and clopidogrel therapy: 2013 update. Clin Pharmacol Ther 94: 317–323.

